# Cell contacts and pericellular matrix in the Xenopus gastrula chordamesoderm

**DOI:** 10.1101/2023.07.05.547782

**Authors:** Olivia Luu, Debanjan Barua, Rudolf Winklbauer

**Affiliations:** Department of Cell and Systems Biology, University of Toronto, Toronto, Canada M5S 3G5; Department of Medical Oncology and Center for Functional Cancer Epigenetics, Dana-Farber Cancer Institute, Boston, MA 02215, USA, and Department of Medicine, Harvard Medical School, Boston, MA 02115, USA

**Keywords:** Xenopus, gastrula, mesoderm, cell adhesion, pericellular matrix, glycocalyx Running title: Cell contacts in chordamesoderm

## Abstract

Convergent extension of the chordamesoderm is the best-examined gastrulation movement in Xenopus. Here we study general features of cell-cell contacts in this tissue by combining depletion of adhesion factors C-cadherin, Syndecan-4, fibronectin, and hyaluronic acid, the analysis of respective contact width spectra and contact angles, and La^3+^ staining of the pericellular matrix. We provide evidence that like in other gastrula tissues, cell-cell adhesion in the chordamesoderm is largely mediated by different types of pericellular matrix. Specific glycocalyx structures previously identified in Xenopus gastrula tissues are absent in chordamesoderm but other contact types like 10-20 nm wide La^3+^ stained structures are present instead. Knockdown of any of the adhesion factors reduces the abundance of cell contacts but not the average relative adhesiveness of the remaining ones: a decrease of adhesiveness at low contact widths is compensated by an increase of contact widths and an increase of adhesiveness proportional to width. From the adhesiveness-width relationship, we derive a model of chordamesoderm cell adhesion that involves the interdigitation of distinct pericellular matrix units. Quantitative description of pericellular matrix deployment suggests that reduced contact abundance upon adhesion factor depletion is due to some contact types becoming non-adhesive and others being lost.

## Introduction

The blastocoel wall of the Xenopus embryo consists of an outer epithelial sheet that coats layers of deep cells. For the epithelial layer, subapical adherens and tight junctions are characteristic (Mueller and Hausen, 1995; Shook and Keller, 2008; Winklbauer, 2020) while deep cell adhesive contacts appear amorphous and are interspersed between non-adhesive interstitial gaps. We recently showed that such contacts can be characterized by their width spectra – the frequency distributions of membrane-membrane distances – and by the modifications of the spectra when adhesion factors are depleted (Barua et al. 2021; Barua and Winklbauer, 2022). From the requirement for factors such as fibronectin (FN) or hyaluronic acid (HA), and from the large widths of contacts we concluded that gastrula cell-cell adhesion is largely mediated by the pericellular matrix (PCM) (Winklbauer, 2019; Barua et al. 2021). In the Xenopus early embryo, a fraction of the PCM contains La^3+^ staining material (LSM) (Johnson, 1977), which can be used to further characterize contact types. Thus, we found that some LSM patches resembled known endothelial glycocalyx variants which we termed glycocalyx I, II and III, and which mediated cell adhesion but occurred also on the free cell surfaces at interstitial gaps (Barua et al. 2021).

Previously, we analyzed ectoderm and prechordal mesoderm, representing different germ layers and morphogenetic behaviors. Whereas ectoderm is stretched during epiboly, prechordal mesoderm performs active, directional cell-on-cell migration (Huang and Winklbauer, 2018). Despite the differences, contact types are largely shared between these tissues (Barua et al. 2021; Barua and Winklbauer, 2022). Here, we analyze cell contacts in the chordamesoderm (CM), which is continuous with the ectoderm at its posterior and the prechordal mesoderm at its anterior end. It elongates by convergent extension, a cell intercalation process driven by protrusive activity at the ends of mediolaterally oriented bipolar cells, and by junction remodelling at antero-posterior cell-cell contacts (Pfister et al. 2016; Shindo and Wallingford, 2014; Weng et al. 2022).

Convergent extension depends on CM cell adhesion via the main cadherin of the Xenopus gastrula, C-cadherin (C-cad) (Pfister et al. 2016; Huebner et al. 2021) and on the extracellular matrix protein FN (Davidson et al. 2006), and we depleted these factors using morpholino antisense oligonucleotides. We also knocked down the small transmembrane/extracellular heparan sulfate proteoglycan Syndecan-4 (Syn-4) (Munoz et al. 2006; Barua et al. 2021), and we impeded the synthesis of the large glycosaminoglycan, HA, by knocking down HA synthases Has1 or Has2 (De Grendele et al. 1996). Combined with the analysis of contact width spectra and La^3+^ staining, the knockdown experiments identified new contact types and showed that the CM contact pattern differs from both ectoderm and prechordal mesoderm. In all morphants the abundance of cell-cell contacts was reduced, but this was not paralleled by reduced average adhesiveness of the remaining contacts: a decrease of adhesiveness at low contact widths was compensated by a dramatic increase of contact widths and an increase of adhesiveness proportional to width. From the latter observation we derive a model of CM cell-cell adhesion which is based on the interdigitation of PCM units from opposite membranes.

## Results

### Abundance of cell contacts but not their adhesiveness is reduced in adhesion factor morphants

At the stage 11 middle gastrula, the blastopore has invaginated to form the archenteron, and the CM has involuted at the blastopore lip to elongate in the animal-vegetal direction between the endodermal archenteron lining and the neural ectoderm (Fig.1A). Depletion of any of four select adhesion factors – C-cad, FN, HA and Syn-4 – by morpholino antisense oligonucleotides affects the CM (Barua et al. 2021). For example, knockdown of C-cad arrests CM involution, elongation, and archenteron formation (Fig.1B), like that of FN (Barua et al. 2021), and the depletion of Syn-4 severely alters cell packing and shape but preserves CM movements (Fig.1C).

**Figure 1.**
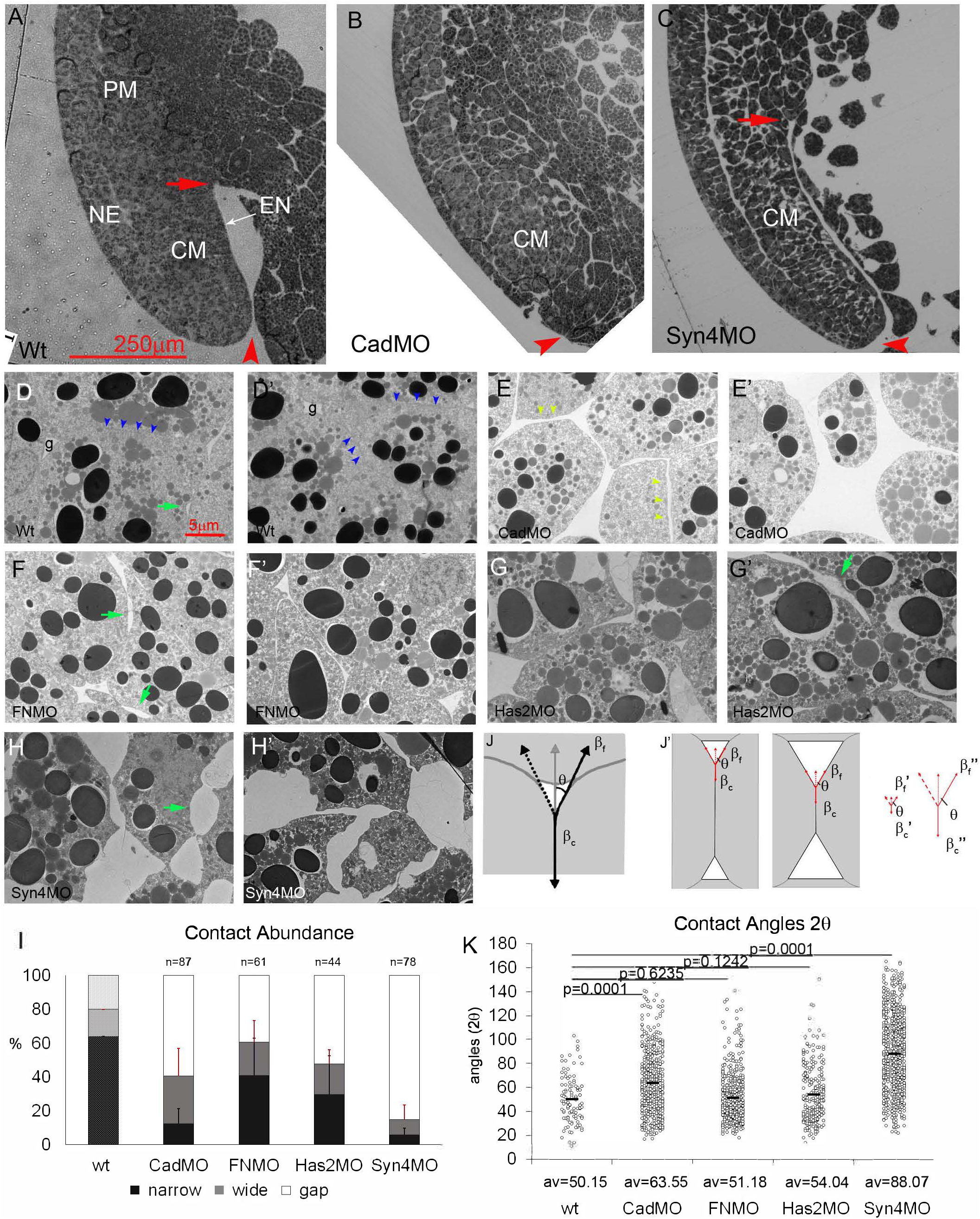
Abundance of cell contacts but not relative adhesiveness is reduced in adhesion factor morphants. (A-C) Dorsal side of normal (A), C-cad depleted (B) and Syn-4 depleted (C) stage 11 gastrulae. CM, chordamesoderm; NE, neural ectoderm; PCM, prechordal mesoderm; EN, suprablastoporal endodermal epithelium; red arrow, tip of archenteron; red arrowhead, position of blastopore. (D-H’) Cell packing in normal (wt) and morphant (MO) stage 11 chordamesoderm. Blue arrowheads, narrow contacts between cells; green arrows, two-sided gaps (“bubbles”) between two cells; light green arrowheads, wide contacts between cells; g, gaps at 3- or 4-cell junctions. (I) Abundances of narrow (<50 nm) and wide (>50 nm) contacts, and interstitial gaps. Bars, standard deviations. (wt) is from Barua et al. 2021. n, number of TEM images analyzed. (J) Tensions at 3-sided gap. For tension βf at free gap surface to balance tension per cell βc at contact interface it must act at a contact angle θ. (J’) Same contact angle θ and thus relative adhesiveness α can combine with large (left) or small (middle) contacts (small or large gaps, respectively). Same θ can be generated by smaller (βf’ and βc’) or larger (βf’’ and βc’’) tensions, provided that their ratio is retained. (K) Each dot represents a measured angle 2θ in normal and morphant CM; av., average.

CM cells are tightly packed, with occasional interstitial gaps at 3-cell junctions and interstitial “bubbles” between two cells (Fig.1D,D’). C-cad knockdown increases the number and size of interstitial gaps, and evenly spaced wide cell-cell contacts become prominent (Fig.1E,E’). FN depletion increases interstitial gaps moderately and widens cell-cell contacts or separates cells (Fig.1F,F’). Knockdown of Has-2 also increases gaps and contact widths (Fig.1G,G’), suggesting an adhesive role of HA. Contact loss is most severe in Syn-4 morphants, where cells remain connected only through thin processes. The cell surface is often concave, generating extensive interstitial space and a serrate cell outline (Fig.1H-H’). Overall, all four factors normally contribute to the dense packing of the CM cells.

To quantify the abundance of adhesive cell contacts (Fig.1I), we measured the lengths of stretches where the membranes of adjacent cells ran parallel until they diverged abruptly at interstitial gaps (Barua et al. 2021). We distinguished between narrow contacts with membrane distances <50 nm, and the remaining, highly variable wide contacts. Non-attached free surfaces are present at interstitial gaps (Barua et al., 2021). In CM cells, 4/5 of the surface is normally engaged in adhesive contacts. Narrow contacts occupy almost 2/3 of the cell surface, and 80-90% of these depend on both Syn-4 and C-cad. Unexpectedly, half of the narrow contacts also require the large HA and FN molecules. As in ectoderm or prechordal mesoderm (Barua et al. 2021), Syn-4 depletion has the strongest effects, suggesting that virtually all narrow contacts and 1/3 of wide contacts require the presence of Syn-4 (Fig.1I). All treatments increase the abundance of non-adhesive gaps.

The abundances of contact widths were collected in 50 nm bins to generate width spectra; changes due to experimental interference are best seen in difference spectra where values of untreated samples are subtracted from experimental ones (Barua et al., 2021). In CM, width abundances decrease sharply after a maximum at <50 nm and vanish beyond 700 nm (Barua et al. 2021) (Fig.S1A). C-cad, FN, Has-2 and Syn-4 depletion strongly reduce abundances of narrower contacts, increase the frequency of wider ones, and generate contacts beyond the normal width range (Fig.S1B-E’). Notably, Has-2 knockdown does not produce the signature glycocalyx I difference spectrum – a distinct decrease at 50-100 nm and increase at <50 nm that is induced by this treatment in prechordal mesoderm or ectoderm (Barua et al. 2021). In fact, compared to the ectoderm (Barua et al. 2021), the 50-100 nm width abundance is lowered in the CM spectrum, reproducing the glycocalyx I depletion signature in the CM-ectoderm difference spectrum (Fig.S1A’) and suggesting a lack of this structure in the CM.

The adhesive strength of contacts depends on the difference in surface energy between free and contacting cell surfaces, β_f_ – β_c_, i.e., on how much tension at contacts is reduced relative to the free surface, for example by the release of adhesion factor binding energy. The ratio of the tensions is β_c_/β_f_ = cosθ with θ the contact angle between adjacent cells. The less the tension is reduced at contacts, the smaller the adhesion strength and the smaller θ (Fig.1J) (Winklbauer, 2015; David et al. 2014). However, both tensions being altered proportionally would change adhesion strength without affecting θ (Fig.1J’). In the CM, angles remain the same upon FN- and Has-2 knockdown and are increased by C-cad and Syn-4 depletion (Fig.1K). Thus, while adhesive contact abundances are diminished in morphants, the relative reduction of tension at the residual contacts – their relative adhesiveness – is not decreased.

### La^3+^ staining and adhesion factor depletion identify chordamesodermal contact types

La^3+^ stains sections of the adhesive contacts, and together with the knockdown of adhesion factors, this can be used to identify contact types. In the ectoderm, isolated LSM plaques occur in contacts and on the surface of interstitial gaps (Fig.2A) (Barua et al. 2021). Cells in the more densely packed CM are often outlined by delicate lines of La^3+^ staining, which are occasionally interrupted, meet at gap-less 3-cell junctions (Fig.2B), can be as narrow as 10 nm (Fig.2C), or, in triple-layered contacts, resolve into two parallel lines with or without small LSM dots between (Fig.2E). Other contacts and rare gaps at 3-cell junctions were unlabeled (Fig.2D). No bush-like glycocalyx II or brush-like glycocalyx III structures (Barua et al. 2021) were observed.

**Figure 2.**
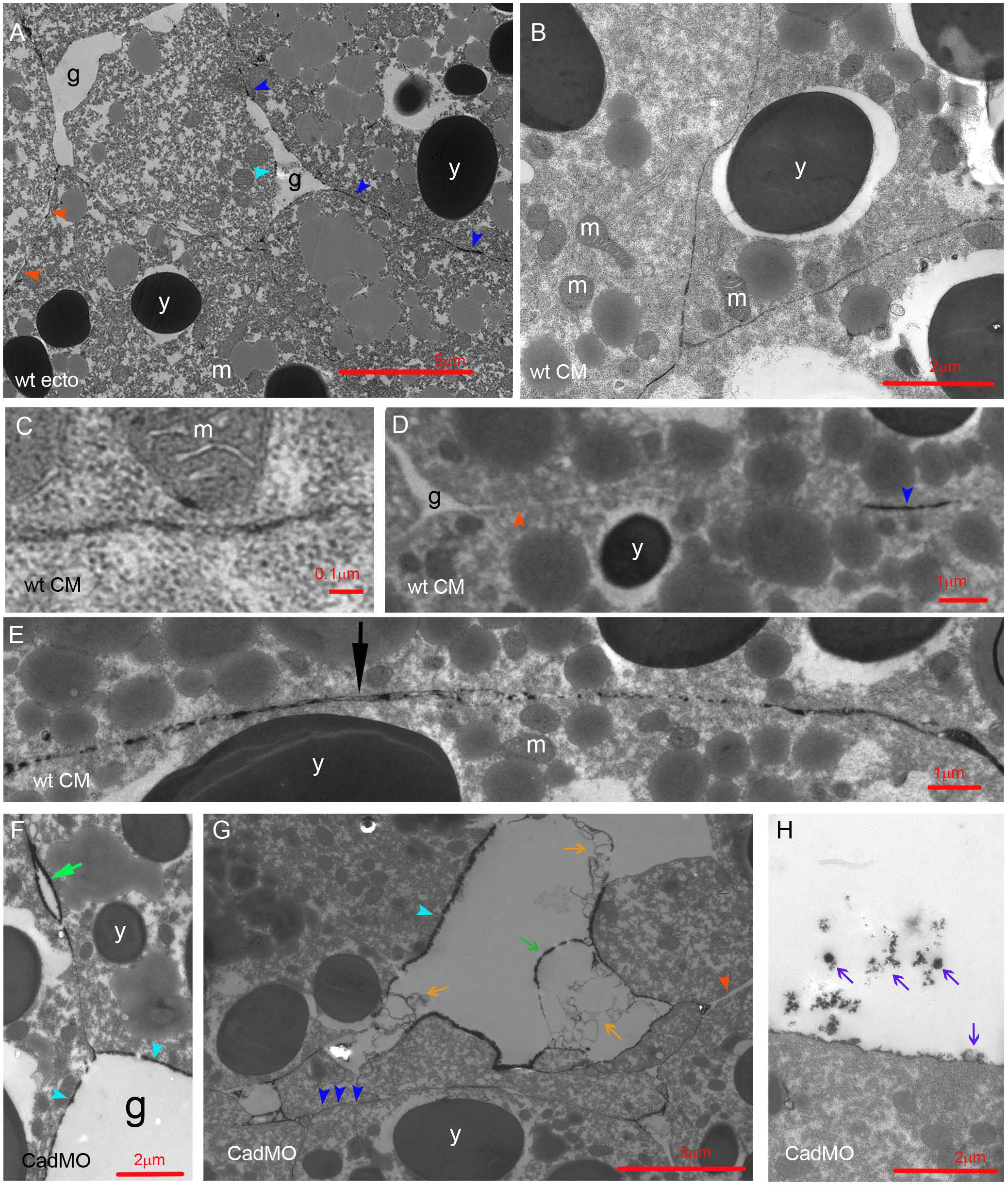
LSM in normal and C-cad depleted CM. (A) LSM in ectoderm, for comparison. (B-E) LSM in normal CM. (F-H) C-cad morphant CM. g, interstitial gaps; y, yolk platelets; m, mitochondria; dark blue arrowheads, LSM plaques in contacts; light blue arrowheads, plaques on gap surfaces; red arrowheads, LSM-free contacts; black arrow, triple-layered contact; light green arrow, bubble; dark green arrow, shed plaque; orange arrows, lightly stained shed ribbons; purple arrows, darkly stained shed LSM material.

Upon C-cad depletion, LSM becomes concentrated in plaques on the surfaces of interstitial gaps (Fig.2F). FN morphants show a similar pattern (Fig.3A,C), and moreover, parallel lines of strong staining can enclose a less stained central layer in contacts (Fig.3C,D). Depletion of HA generates long unlabeled contacts (Fig.3E) or breaks LSM lines up into short plaques or rows of droplets that sit with a broad base on one cell surface and touch with their apex the opposite cell (Fig.3F). In Syn-4 morphants, cell surfaces are mostly devoid of La^3+^ staining (Fig.3G), but 10-20 nm LSM contacts are preserved (Fig.3H), and some surfaces are coated with small, sparse LSM dots (Fig.3I). Three types of LSM shedding occur. In gaps of C-cad or FN morphants, complete plaques and long, faintly stained ribbons detach (Figs.2G, 3B); groups of small, dense LSM flakes are also shed (Figs.2H, 3B); and in Syn-4 morphants, hairballs of fine fibrils (Fig.3J) consist probably of HA (Barua et al. 2021). Thus, as in ectoderm or prechordal mesoderm, Syn-4 is the main contributor to LSM formation. Syn-4 and HA seem to generate extended LSM plaques, which are prone to shedding though in the absence of C-cad or FN.

**Figure 3.**
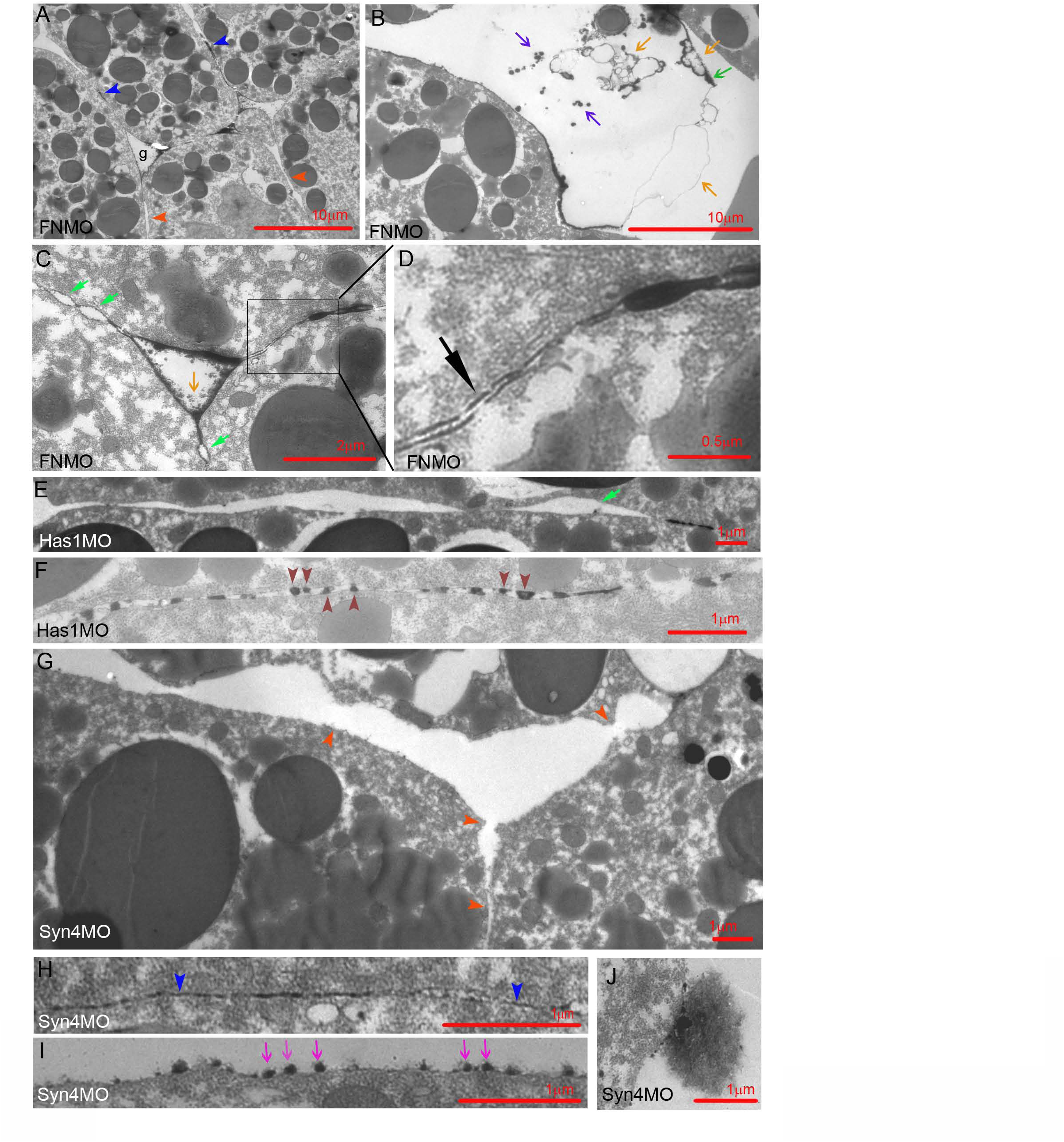
LSM in FN, Has1 and Syn-4 depleted CM. Dark red arrowheads, semi-drop-like LSM sitting on upper or lower membrane; dark blue arrowheads, LSM plaques in contacts; red arrowheads, LSM-free contacts; black arrow, triple-layered contact; light green arrow, bubble; dark green arrow, shed plaque; orange arrows, lightly stained shed ribbons; purple arrows, darkly stained shed LSM; magenta arrows, LSM dots.

LSM-filled contacts were further characterized by their width spectra. In ectoderm, width abundances decrease from a maximum at 20-30 nm (Fig.4A). In the CM (Fig.4B,B’), widths decrease first gradually from a maximum at 10-20 nm, then abruptly at 60 nm, and abundances remain low in the 60-120 nm range which in ectoderm harbors glycocalyx I, consistent with this glycocalyx type being diminished in the CM. A new CM peak at 10-20 nm is apparent in the CM-ectoderm difference spectrum (Fig.4B’), and C-cad-MO (Fig.4C, C’) and Syn-4-MO (Fig.4F,F’) spectra confirm the subdivision of narrow LSM contacts into C-cad/Syn-4-independent 10-20 nm and -dependent 20-50 nm contacts. In FN and Has1 morphants, all LSM contacts <50 nm are strongly diminished, and 50-100 nm contacts increased. HA and FN are particularly required in 10-20 nm contacts (Fig.4D-E’). In C-cad and Syn-4 morphants, triple-layered contacts (see Fig.2E) are absent, and in Has-1 and FN morphants, their peak is shifted from 30 to 70 nm (Fig.S2A-C). Thus, C-cad and Syn-4 are essential for these contacts while HA and FN control their width.

**Figure 4.**
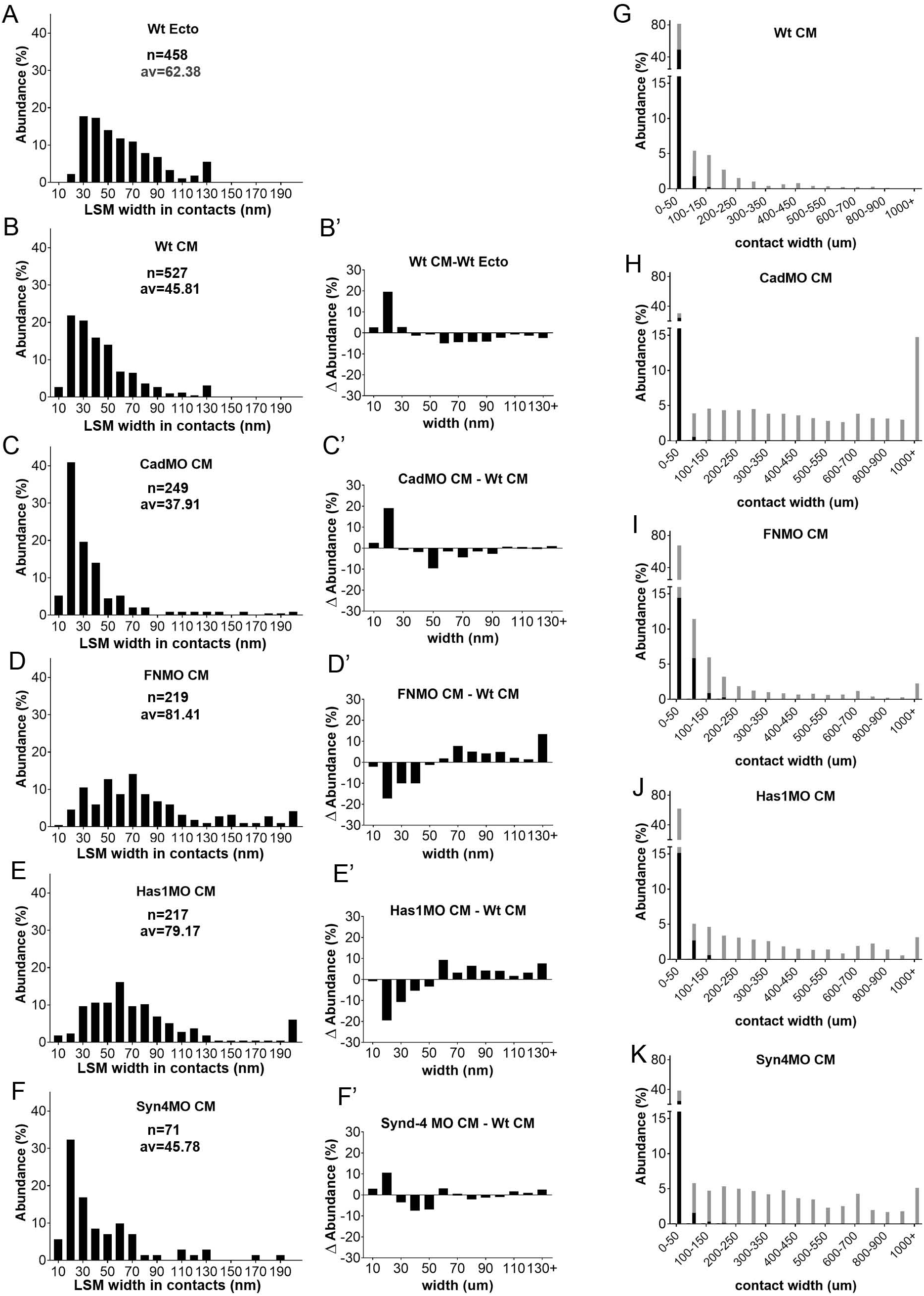
LSM widths in contacts. (A-F) Width frequency distributions in ectoderm (A) for comparison, in CM (B) and in various CM morphants (C-F). n, number of width measurements from 18, 4, 6, 8, 3 TEM images, respectively; av., average. (B’-F’) Corresponding difference (ΔAbundance) spectra comparing width distributions of CM to ectoderm (B’), and of morphants to normal CM (C’-F’). (G-K) Comparison of widths of LSM-containing (black parts of bars) and LSM-free (grey parts of bars) contacts, using the data from (B-F) and Figure S1.

A large part of the PCM is not stained by La^3+^ and respective contacts are less well defined (Fig.4G-K). Almost half of narrow <50 nm contacts are unlabeled in the CM. They are essentially removed by C-cad and Syn-4 knockdown while wider than normal contacts appear, consistent with a widening of the original contacts. In FN and Has1 morphants, unlabeled contacts are little affected. Thus, narrow C-cad- and Syn-4-dependent unlabeled PCM contacts form part of the CM contact complement. Of note, LSM is restricted to widths below 200 nm, leaving the wider contacts, predominant in morphants, unlabeled.

### PCM distribution in contacts and in non-adhesive gaps

LSM is typically more prominent in non-adhesive gaps than in contacts (Fig.5A-F). Thick layers of LSM can accumulate symmetrically on neighboring sides of a gap in C-cad, FN and Has morphants while the width of an adjoining LSM contact is much thinner than the height of the gap surface LSM (Fig.5A-D). Examples from the more frequent gaps in prechordal mesoderm show that this is also true for normal tissue (Fig.5.E,F). Notably, when the LSM forms distinguishable units, these seem to interdigitate in contacts (e.g. Figs.5B,E; see also Barua et al. 2021) to generate contacts only as wide as the units are high in each single layer. Compared quantitatively to LSM contact widths in normal CM (see Fig.4), even single LSM layers are higher in gaps in the morphants (Fig.5G-J), and the respective frequency distributions indicate 2.2-fold, 2.8-fold, 4.0-fold and 1.4-fold increases in gaps of C-cad, FN, Has1 and Syn-4 morphants, respectively. Gap-contact difference spectra confirm reduced abundances of narrow LSM layers and an increase of thicker layers, consistent with the width of contacts not being derived from adding up the heights of LSM layers in gaps.

**Figure 5.**
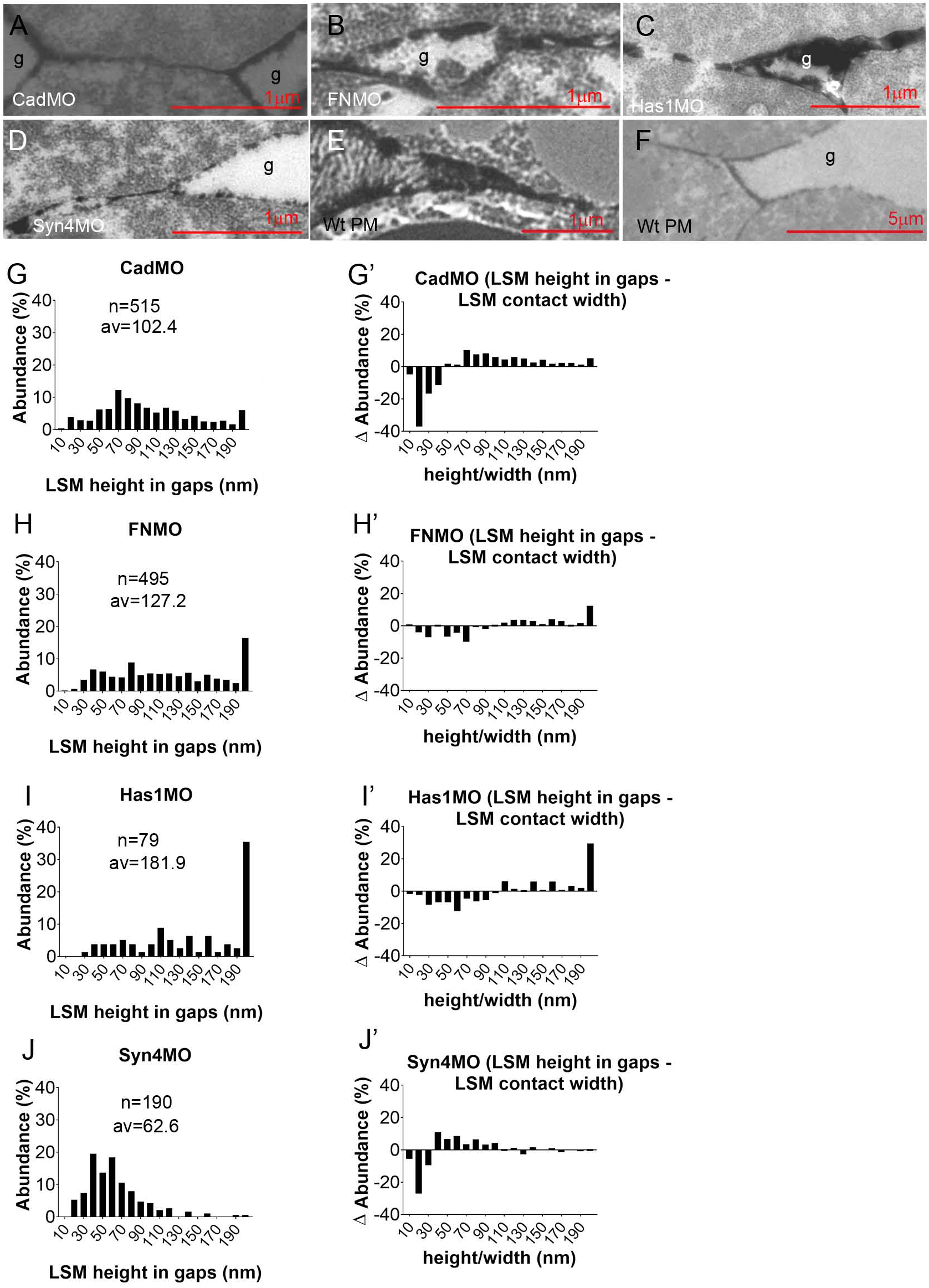
LSM in gaps. (A-F) Examples of LSM at transitions from gaps (g) to cell-cell contacts. PM, prechordal mesoderm. (G-J) Frequency distributions of LSM height in gaps (corresponding to LSM width in contacts). n, number of measurements from 5, 6, 3, 7 TEM images, respectively; av., average. (G’-J’) Difference (ΔAbundance) spectra corresponding to (G-J), comparing LSM height in gaps to LSM width in contacts.

In CM contacts, the lengths of continuous LSM, non-stained, or triple-layered stretches are similar (Fig.6A), and similar to those of LSM plaques in prechordal mesoderm (Barua et al. 2021). In morphants, LSM stretches are lengthened or shortened moderately while the lengths of non-stained stretches are always increased, in CadMO embryos by 10-fold (Fig.6B-E). Random removal of some LSM plaques interspersed in non-labeled contact stretches would cause this effect. LSM plaque length in gaps is comparable to that in contacts in C-cad and Syn-4 morphants and increased in FN and Has1 morphants (Fig.6B’-E’). The lengths of unlabeled stretches in gaps (Fig.6B’-E’) suggest also random removal or addition of LSM stretches in non-labeled regions.

**Figure 6.**
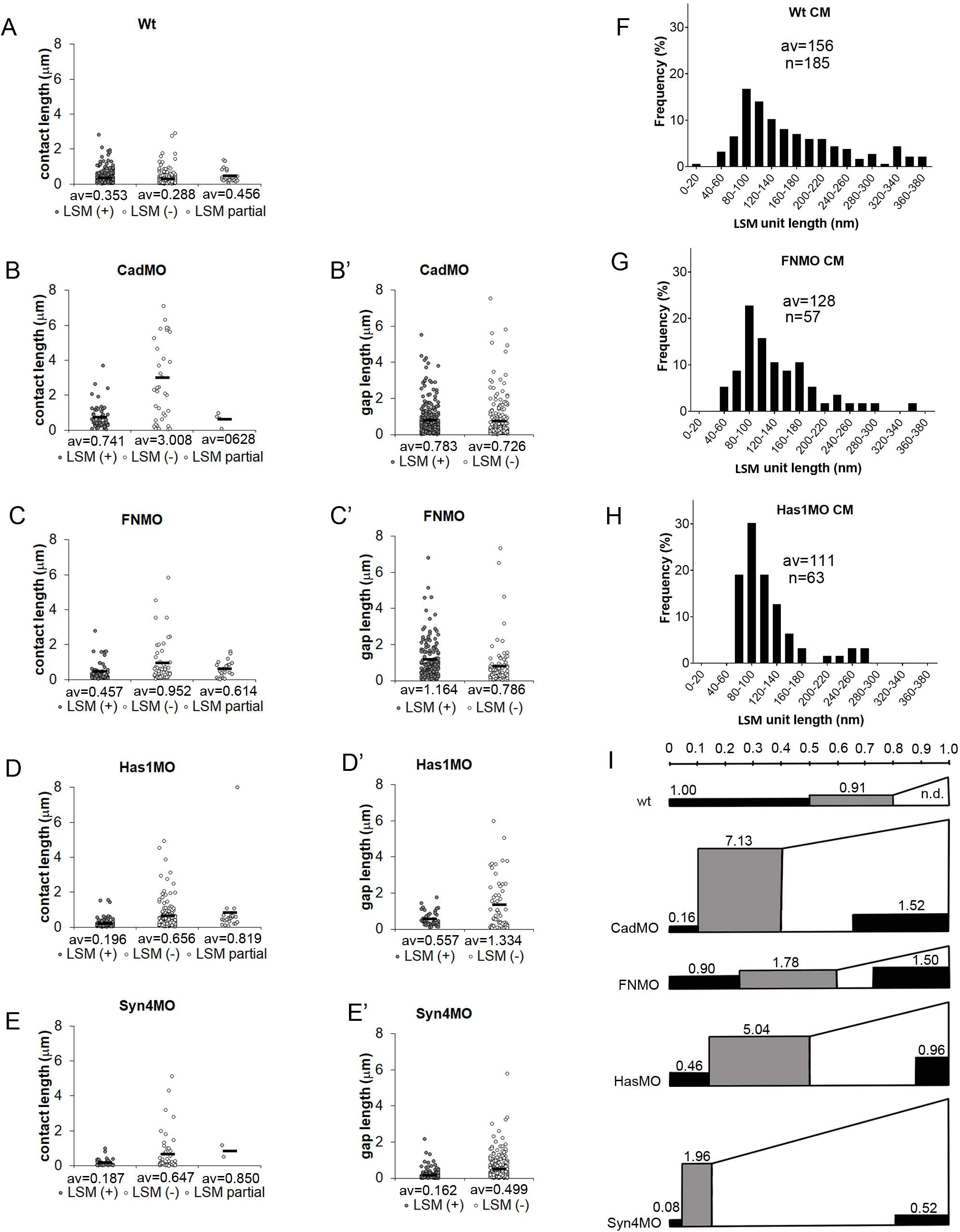
Lengths of PCM stretches. (A-E) Lengths of continuous stretches of LSM-containing (LSM(+) contacts (plaques), and of LSM-free (LSM(-) and triple-layered (LSM partial) contacts. (B’-E’) Lengths of continuous stretches of LSM (plaques) (LSM(+)) on gap surfaces or LSM-free surfaces (LSM(-)). Data from 4, 5, 9, 8, 6 TEM images, respectively; av., averages. (F-H) Lengths of shortest LSM units discernible in normal CM contacts (F), and in contacts and gaps of morphants (G,H). n, number of unit LSM structures measured, from 4, 7, 5 TEM images, respectively. (I) Summary diagram integrating LSM (black areas) height and length data for contacts (left, rectangular) and gaps (right, wedged), and non-labeled contact height and length data for contacts (grey, rectangular). Area size relative to LSM in normal CM contacts (1.00) is indicated. Scale on top, relative average lengths of contact types. Height of rectangles, relative average widths of respective structures. Areas are proportional to respective PCM volumes.

To see whether large plaques could be composed of smaller units, we measured the lengths of the smallest discrete LSM structures in normal CM contacts (e.g. Fig.2E), in contacts and gaps of FN morphants (e.g. Fig.5B), and the LSM droplets in HA depleted CM (e.g. Fig.3F). We consistently found a peak at 100 nm with a slow decrease towards higher values (Fig.6F-H). In the plaque size distribution of CM (Fig.6A), these unit structures would occupy the lower margin. This is consistent with larger plaques being assembled from small units, and combinations of different LSM and non-LSM units generating patterns of alternating contact types.

The effects of adhesion factor depletion on PCM distribution are summarized in (Fig. 6I). In CM, more than half of adhesive contact length is La^3+^-stained. In morphants, overall contact length is reduced and LSM contacts shrink disproportionally, from half of the total cell surface in normal CM to ¼ or 1/5^th^ in FN, C-cad and Has2 morphants, and less than 1/10^th^ upon Syn-4 depletion. Non-labeled contacts remain at normal lengths except in Syn-4 morphants. In randomly sliced samples, volumes of object are proportional to their averaged cross-section areas, and for the volume of LSM or unlabeled PCM in contacts, relative contact length was multiplied by relative width (Fig.6I). C-cad or Syn-4 depletion reduce the contact volume of LSM by an order of magnitude. HA and FN depletion combine shortening of LSM contacts with their widening. Non-labeled contacts, although shorter, are wider in normal CM and their volume equals that of the LSM contacts (Fig.6I). In C-cad morphants, width is dramatically increased, and the volume of non-labeled contact augmented by almost 8-fold. Width increases lead also to 2–5.5-fold higher contact volumes in the other morphants (Fig.6I). The volumes of LSM and non-labeled contacts combined increase between 1.4- and 3.8-fold in the morphants, except for Syn-4 knockdown where the total is unchanged.

In gaps (Fig.6I), more than half of the cell surface is coated with LSM in FN and C-cad morphants, and almost a quarter in HA or Syn-4 depleted CM. LSM layer thickness is increased compared to contacts, and together with the increase in overall gap size in all morphants, this amounts to a large LSM volume in gaps which dominates the total volume of LSM. In FN morphants, total LSM volume exceeds that of untreated CM by 2.5-fold, with the excess being accumulated in the gaps. In C-cad and Has1 morphants, a reduced LSM volume in contacts combines with increased LSM in gaps, as if material were redistributed with only a modest increase in total volume. In Syn-4 morphants total LSM volume is reduced by half, suggesting that Syn-4 promotes LSM deposition or is itself a main part of the LSM (Barua et al. 2021). The amount of non-labeled PCM in gaps remains unknown, but even if it were completely absent, adhesion factor depletion causes a considerable increase of total PCM volume. This could be due to the overproduction of PCM material, to the swelling of existing PCM, or both.

### Relative adhesiveness is a function of contact width

Contact angles and contact width are both spread over a wide range in normal and morphant CM, and we asked whether the two parameters are correlated. Importantly, the contact angle θ between cells is related to the dimensionless relative adhesion strength α, i.e. at gaps to the adhesion strength σ_i_ = β_f_ – β_c_ normalized to the tension at free surfaces, α = σ_i_/β_f_ = 1 – β_c_/β_f_ = 1 – cosθ (Figs.1J, S5) with 0 (no adhesion) ≤ α ≤ 1 (maximal adhesion) (Winklbauer, 2015; David et al. 2014). Overall, α increases with width w in normal and in morphant CM (Fig.7A-E). In respective scatter plots, α values are concentrated above a lower boundary but become more dispersed farther above and with increasing w. In normal CM, most α values reside at w < 250 nm, and their average in this range is significantly lower in C-cad, FN, and Has morphants compared to normal CM (Table S1). However, for w > 250 nm, α is strongly increased, and the expanded width ranges in the morphants allow to compensate their reduced basic adhesiveness such that overall relative adhesion strength remains normally high. Syn-4 morphants differ, showing increased average α at both width ranges. The α–w relationship is conveniently analyzed for the lower boundary of α values. We constructed it by selecting the data points on the enveloping curve, proceeding from one such point to the next point which is higher but does not require moving lower again subsequently. This was repeated up to where points became sparse (Fig.7A’-E’). A linear regression line was then fitted through the enveloping points as α = (Δα/Δw)w + α_0_ with Δα/Δw the slope of the line and α_0_ its extrapolated value at w = 0 (Fig.7A-E’). These parameters were similarly decreased in C-cad, FN and Has morphants compared to normal CM. Syn-4 morphants had increased values (see Table S3).

**Figure 7.**
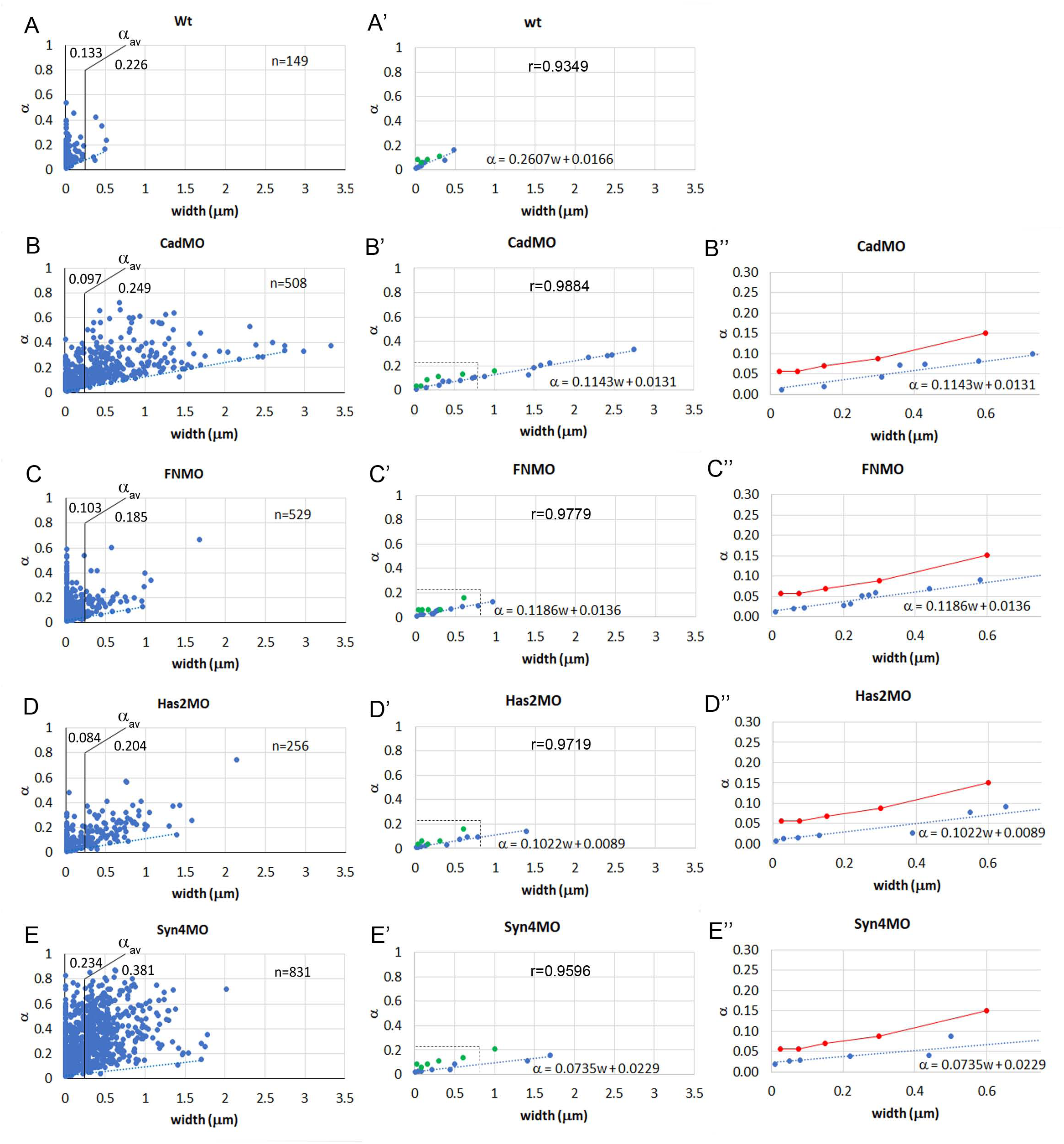
Relationship between relative adhesiveness and contact width. (A-E) Relative adhesiveness α was determined from contact angles and plotted as a function of contact width w for each gap-contact transition. The average value of α, α_av_, is indicated for w smaller or larger than 250 nm (see Table S1). n, number of transitions measured in 26, 86, 59, 41, 80 TEM images, respectively. (A’-E’) A linear regression line (dotted blue line) was fit to the lowest values in each plot (blue dots). The slope Δα/Δw and the α-axis intercept α_0_ of the regression line α = (Δα/Δw)w + α_0_ are indicated. r, correlation coefficient for regression line. Green dots, values for α frequency peaks (see Fig.8 and Table S2). (B’’-E’’) Higher magnification of (B’-E’) focussing on small w. α frequency peak data in (B’-E’) were pooled (red dots and lines) (see Table S2) and compared to lower-boundary regression lines (blue).

To examine the α–w relationship above the lower boundary, we determined the frequency distributions of α values for consecutive width brackets. Overall, the distributions are skewed with a first peak near the lower boundary and a long tail tapering off irregularly at higher values (Fig.8A-E; Fig.S3). Except for Syn-4 morphants, this first peak is usually the main peak of the distribution. It is not shifting noticeably to higher values over the first two width brackets, centered around 25 and 75 nm (Fig.S3), which were thus combined in Fig.8A-E. As distributions shift to higher α values with widths above 200 nm, minor peaks or shoulders appear or become more prominent, and differences between treatments more obvious (Fig.8A-E). Particularly, in Syn-4 morphants, frequencies are spread out almost evenly over a large α range (Fig.8E).

**Figure 8.**
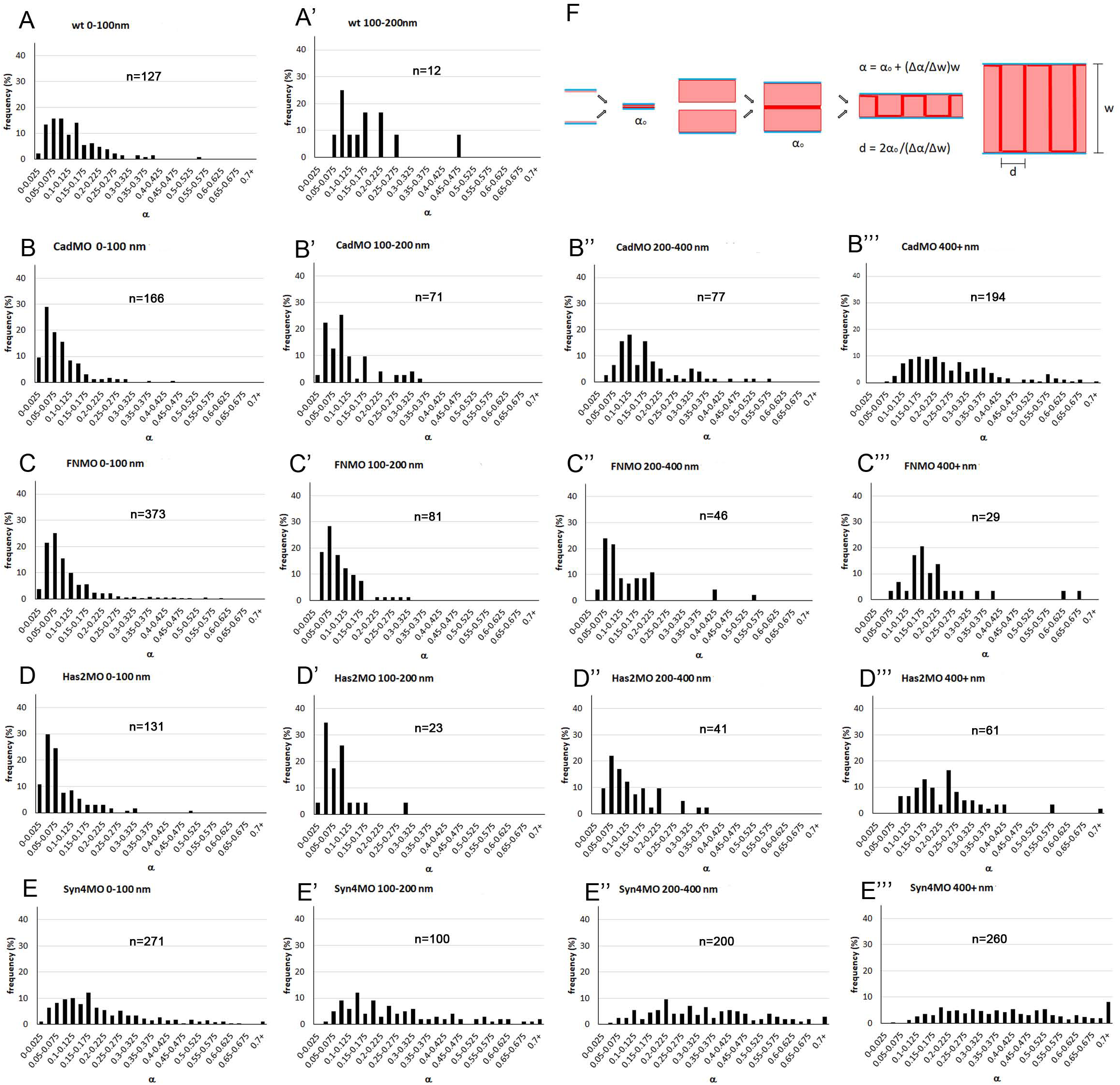
Frequency distributions of α values at different widths. (A-E’’’) Different treatments are arranged vertically and width brackets horizontally, as indicated on top of each diagram. n, number of α – w data points. Additional width brackets are shown in Figure S3. (F) Model of cell-cell adhesion by PCM interdigitation. Adhesion between two thin or thick PCM layers (light red) on cell membranes (blue) occurs in principle through a narrow interaction zone (deep red), yielding in each case the basic adhesiveness α_0_ (left). Interdigitation of the two apposed PCMs corresponds to a folding of the interaction surface which increases linearly with PCM height w at constant interdigitation distance d (right).

The frequency of α values declines rapidly with increasing w (Fig.7A-E), and with the heavily skewed α frequency distributions (Fig.8), large α values disappear preferentially at higher w, artificially lowering the nominal average of α. Instead, the first peak of the α frequency distributions can be followed to analyze the α–w relationship above the lower boundary (Fig7A’-E’’). Only few points, at low resolution, are obtained (Fig.7A’-E’), but the values for all morphants are similar (Table S2) and are thus pooled. The line connecting the averages increases linearly for w > 100 nm, but at a steeper slope than the lower-boundary line. For smaller widths, it seems to remain constant, in contrast to the lower boundary (Fig.7B’’-E’’).

Contact-gap transitions are sectioned at randomly oriented planes and distortion of contact angles broadens their distribution, letting narrow angles appear even narrower and wide angles even wider (Barua et al. 2017). The variation of α values is broadened accordingly by the factor k, and we estimated k ≈ 4/3 (see Materials and Methods). Correction by this factor tilts lower boundary and peak lines upward (Fig.S4). The effect should be the same for all treatment conditions, and the span of the different α frequency distributions is also of the same order of magnitude. It is the range of contact widths w that differs several-fold between treatments, and together with α ∼ w, this determines the overall relative adhesiveness of normal and morphant CM.

## Discussion

### Cell-cell contact types in the chordamesoderm

In terms of cell packing density, in vitro cell motility, and contact spectra, the CM resembles the ectoderm more than the prechordal mesoderm (Wacker et al. 1998; Barua et al. 2021). However, ectoderm and prechordal mesoderm share contact types while the CM, located between and linking these two regions, is different. In prechordal mesoderm and ectoderm, a glycocalyx I is identified via its difference spectrum characteristics, and glycocalyx II and III are recognized by their morphology (Barua et al. 2021). None of these structures are seen in the CM. Instead, a 30 nm wide, triple-layered contact type with a non-labeled layer between two LSM sheets is present in the CM. Another, 10-20 nm wide LSM contact in the CM is narrower than adherens junctions and insensitive to C-cad depletion but depends on the large HA and FN molecules. Despite their size, these factors can reside in narrow spaces. CNS synapses 20 nm wide harbor HA and its CD44 receptor (Roszkowska et al. 2016; Wilson and Litwa, 2021), and the string-like FN protein likewise occurs in synaptic clefts (Thalhammer and Cingolani, 2014). HA chains thousands of nm long occupy large volumes when randomly coiled, but when attached to a surface can form 0.3 nm thin layers (Jacoboni et al. 1999). Thus, HA and FN could directly build the 10-20 nm contacts. Non-LSM contacts are prominent in the gastrula but due to the lack of respective staining are less well characterized. In the CM, some narrow < 50 nm non-LSM contacts depend on C-cad, consistent with the presence of adherens junctions (Müller and Hausen, 1995).

### Pleiotropic and polygenic control of contact complexes by adhesion factors

The CM results confirm our previous findings that adhesion factors are pleiotropic, each affecting various contact types, while contact types in turn are each controlled by several adhesion factors (Barua et al. 2021; Barua and Winklbauer, 2022). As an example for pleiotropic functions, HA acts together with Syn-4 and other factors in ectoderm and prechordal mesoderm to build a 50-130 nm wide glycocalyx I, and in prechordal mesoderm a micron-wide brush-like glycocalyx III (Barua et al. 2021). In the CM, HA supports instead extremely narrow 10-20 nm LSM contacts. On the other hand, glycocalyx I is the prime example for a structure that depends on multiple adhesion factors, in the prechordal mesoderm at least on HA, Syn-4, Glypican-4, PAPC, ephrinB3 and EphB4 (Barua et al. 2021; Barua and Winklbauer, 2022). The CM provides additional examples. Among only 4 factors tested, the 10-20 nm contacts require HA and FN, some 30-50 nm contacts C-cad and Syn-4, and triple-layered contacts all four factors.

The picture emerges that different adhesion factors, in variable combinations, build a mosaic of adhesive contact complexes whose molecular and mechanical details are yet to be studied. Depletion of a factor often modifies a contact. For example, upon depletion of HA, 10-20 nm LSM contacts disappear and wider, beaded LSM contacts appear, suggesting a change in width and structure, but not in adhesive role. Likewise, triple layered LSM contacts become wider in FN morphants, and non-LSM contacts apparently in all morphants. Modification, not dismantling was also observed in endothelial glycocalyx when HA or heparan sulfate were enzymatically removed (O’Callaghan et al. 2011). Modification of a contact type could include becoming non-adhesive. Thus, the apparent redistribution of LSM from contacts to gaps could be due to the accumulation in gaps of LSM rendered non-adhesive. The overall reduction of LSM upon Syn-4 depletion and the absence of triple layered contacts in C-cad and Syn-4 morphants suggest that contact types can also be lost. Quantitatively similar α–w curves in different morphants and across contact widths ranges suggest similar adhesiveness of the different residual adhesion types. Predominantly non-specific adhesion between PCMs by van der Waals forces, hydrogen bonds, Ca^2+^ bridges, or chain entanglement (Han et al. 2008; Boettiger and Wehrle-Haller, 2010; Vilanova et al. 2016; Even et al. 2017; Cao and Forrest, 2019) would contribute to this, while specific interactions of adhesion factors could play mostly structural roles in the PCM.

### Adhesion strength in the chordamesoderm

The average relative adhesiveness α at gaps in normal and in different adhesion factor-depleted CM is not correlated with contact abundance. To understand this unexpected result, we ask how relative adhesiveness α is related to a measure of absolute adhesion strength, the tissue surface tension σ, which has been determined for Xenopus gastrula tissues including the CM (Ninomiya et al. 2012; David et al. 2014; Kashkooli et al. 2021; Shook et al. 2022). It corresponds to the difference between the tensions at the free tissue surface, β, and at cell contacts, β^c^. β is essentially the tension generated by the cell cortex, and β_c_ is determined by a cadherin-dependent downregulation of this cortical tension to a within-tissue level β_f_ and an oppositely oriented adhesion tension Γ/2, the binding energy released per unit area and per cell upon adhesion (Fig.S5) (Manning et al. 2010; Winklbauer, 2015). Thus, σ = β – (β_f_ – Γ/2), with β – β_f_ being large compared to Γ/2 in gastrula tissues (David et al. 2014; Barua et al. 2017). The contact angle θ_s_ at the tissue surface is given by cosθ_s_ = (β_f_ – Γ/2)/β (Fig.S5). At the free surface of interstitial gaps, β_f_ is known to be preserved (Barua et al. 2017). Tension at contacts is again β_f_ – Γ/2, and adhesion strength between cells attaching and detaching within the tissue is σ_i_ = β_f_ – (β_f_ – Γ/2) = Γ/2. Tension equilibrium requires a contact angle θ much smaller than that at the tissue surface, according to cosθ = (β_f_ – Γ/2)/β_f_ (Fig.S5A).

For the strength of C-cad mediated adhesion, quantitative data are available. Depleting C-cad reduces adhesion strength σ in gastrula tissues by about half: β is less reduced at contacts, and thus the difference σ between increased β_c_ and unchanged β is smaller (David et al. 2014). At unaltered adhesion tension Γ/2, this decreases contact angles θ at gaps and thus α (Fig.S5B), consistent with the diminished average α in the < 250 nm width range in C-cad morphants. Most contacts are much wider though in these morphants, and as α increases with w, the average α for w > 250 nm is much higher and the simplest explanation for this effect is that Γ/2 is also a linear function of w. The overall increase of Γ/2 upon C-cad depletion tends to compensate the increase in β_f_ (Fig.S5C). Quantitatively, from our previous data for β_c_ in normal CM (0.06 mJ/m^2^) (David et al. 2014), we calculate a β_f_ of 0.07 mJ/m^2^ and Γ/2 of 0.01 mJ/m^2^, values which are almost identical for the ectoderm (Barua et al. 2017). Assuming a similar increase of β_c_ as in the ectoderm upon C-cad depletion, to 0.15 mJ/m^2^, β_f_ would be 0.19 mJ/m^2^ and Γ/2, 0.04 mJ/m^2^, i.e. β_f_ increased by 2.5-fold and Γ/2 by 4-fold, suggesting overcompensation, as apparent from the contact angles in Fig.1K. A similarly raised, by then unexplained, Γ/2 after C-cad depletion had also been found for the ectoderm (Barua et al. 2017).

In summary, at gaps the effect of C-cad knockdown on cortical tensions is compensated by an increase in average adhesion tension Γ/2 due to the widening of contacts in morphants and a proportional increase of adhesiveness. Adhesion strength changes in FN and Has morphants have yet to be examined. The observed initial reduction of adhesiveness at a given width could in principle occur as in C-cad morphants (e.g. Stevens et al. 2023) but could also be due to reduced adhesion tension. If the observed increase in contact PCM volume was due to swelling and accompanied by a decrease in PCM density, less interaction between binding sites would take place per PCM-PCM interface area and less binding energy would be released. Adhesion in Syn-4 depleted CM appears very different and may require different concepts for its analysis.

### A mechanism of pericellular matrix-based cell-cell adhesion

To mediate cell-cell adhesion, PCM surfaces have been proposed to interact molecularly in a narrow contact zone by the limited interdiffusion of chain ends and the exchange of non-covalent binding interactions (Winklbauer, 2019). With only a thin slice at the surface of PCMs involved, adhesiveness would be independent of total PCM thickness and hence of contact width (Fig.8F). For w < 100 nm, this is indeed seen for the line tracing α frequency peaks, but for this line at w > 100 nm and for the whole low-boundary line, the linear increase of α with w implies that adhesive interaction increases with the thickness of the PCM, most simply achieved by the interpenetration of the PCMs of two cells. Complete interpenetration of the glycocalyx layers has been observed at erythrocyte-macrophage contacts (Soler et al. 1998). In the gastrula, the observed change in LSM layer thickness at transitions between contacts and gaps is consistent with such a process.

Interpenetration of coherent PCM meshworks, molecule by molecule, over hundreds of nanometers, seems unlikely. To estimate the diameters of putative interpenetrating units in the contact plane, we assume that the units adhere on all their sides, and that the slope (Δα/Δw) of the α–w curve is thus proportional to the density of unit-unit contacts, i.e. to twice the inverse of the diameter d of a unit, Δα/Δw ∼ 2/d. Further, the slope should be proportional to the basic PCM-PCM adhesiveness α_0_, as seen in the absence of interpenetration, and thus Δα/Δw = 2α_0_/d. For calculations the lower-boundary regression line provides the best data. Here α → α_0_ for w → 0 (Fig.8F) and with α_0_ and Δα/Δw read off from the α–w curves (Fig.7), d can be calculated as between 130-235 nm, except for Syn-4MO (Table S3). The main peaks from the α distributions provide only few, low resolution data points for α–w curves but between 100 nm and 600 nm the combined data are consistent with a linear increase, in proportion to the lower-boundary line. Lines corrected for contact angle distortion are constructed by proportionally increasing the original lines by a factor k and will thus also give the same d values. Minor peaks in the α frequency distributions may indicate other contact types in a certain width range, with higher basic adhesiveness.

The estimated d values are an order of magnitude larger than the deduced endothelial glycocalyx molecule spacing of ∼ 20 nm but agree with a 100 nm periodicity in that glycocalyx that reflects the size of glycocalyx bushes (Squire et al. 2001; Weinbaum et al. 2007), and with the sizes of minimal LSM units identified here. Interdigitation of such units is apparent in prechordal mesoderm where glycocalyx II bushes from opposite membranes intercalated in antiparallel fashion at the transition between interstitial gap and adhesive contact (Barua et al. 2021). Contacts wider than 200 nm are not labeled with La^3+^ in the CM, but the linear α–w relationship is maintained, predicting that non-labeled PCM also consists of similarly sized structural building blocks, or PCM stubs, which can interdigitate for adhesion. In summary, the stub interdigitation model assumes a thin PCM-PCM interaction zone, determined by short-range molecular interactions, whose surface is increased by interdigitation equivalent to an effective large-scale folding, such that adhesion tension per cell surface area, Γ/2, increases with PCM width (Fig.8F).

Intercalation unit diameter d of Syn-4 morphants differs strongly from that of other morphants, suggesting a unique contact structure. Overall contact and LSM contact abundances, average relative adhesiveness α, and the α value distribution are also different. At the same time, cell shape differs fundamentally. In a single step, by the depletion of Syn-4, a spindly, serrate mesenchymal cell outline is attained which is common in embryonic tissues of other vertebrates (Trelstad et al. 1967; Granholm and Baker, 1970; Batten and Haar, 1979; Singley and Solursh, 1980) but not in amphibian gastrulae. Remarkably, gastrulation movements including the well-studied convergent extension of the CM are not affected by the mid-gastrula stage by this transformation whose cellular basis remains to be elucidated. We did not examine how axis defects were caused by Syn-4 depletion towards the end of gastrulation (Munoz et al. 2006).

Shedding of whole plaques but also of isolated stubs suggests that stubs are capable of reversible lateral adhesion. This raises the question of how the interdigitation of stubs from opposite membranes is favored over their lateral association as plaques on the same membrane. Possibly, their antiparallel lateral attachment during interdigitation releases more binding energy than parallel contact. Being linked to the cortical cytoskeleton (Weinbaum et al. 2007), stubs within plaques could also be actively separated to facilitate interdigitation, regulating the initiation of adhesion. Conversely, active lateral compression in existing contacts could promote de-adhesion and cell separation. Generally, controlled deployment of PCM materials at contacts and in gaps and regulation of PCM height and density may determine contact abundance and adhesion, and involve loops of cell signaling originating from and aiming back at the PCM. Upon adhesion factor depletion, some contacts are modified but remain adhesive. Others become non-adhesive and still others may disappear, thus lowering contact abundance.

## Materials and Methods

### Embryo manipulations

Adult *Xenopus laevis* were maintained in accordance with University of Toronto Animal Use Protocol (20011765). Eggs were fertilized in-vitro, de-jellied using 2% cysteine in 1/10 Modified Barth’s Solution (MBS; 88 nM NaCl, 1 mM KCl, 2.4 mM NaHCO_3_, 0.82 mM MgSO_4_, 0.33mM Ca(NO_3_)_2_, 0.41 mM CaCl_2_, 10 mM Hepes (+NaOH), 1% streptomycin, 1% penicillin (pH 7.4) and kept in 1/10 MBS until stage 11. Morpholino antisense oligonucleotides (Gene Tools) were injected at the two-cell stage in 4% ficoll solution and embryos were incubated in 1/10 Modified Barth’s Solution at 15^0^C until stage 11. The following previously characterized morpholinos were used, at the efficiencies as percent reduction of protein levels indicated.

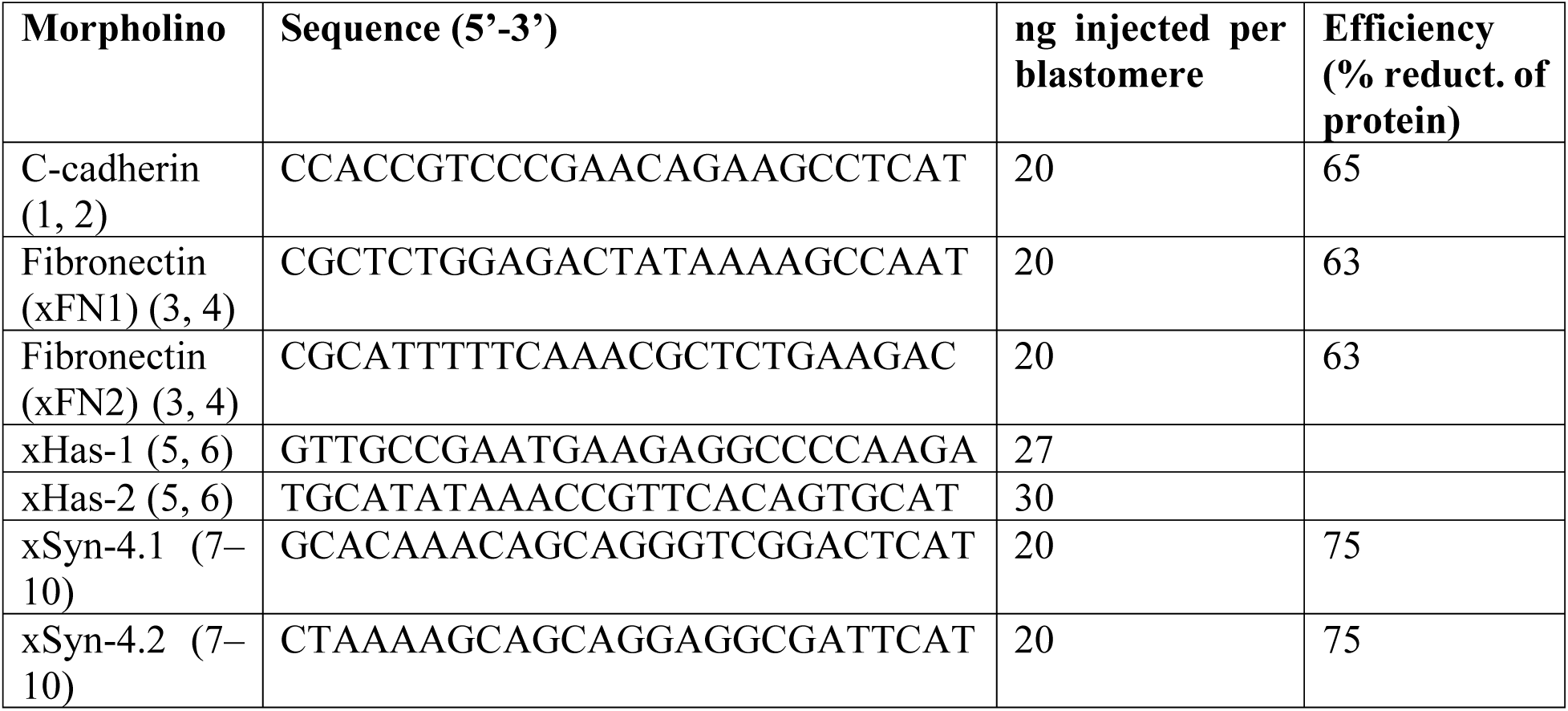

References: (1) Ninomiya et al. (2012); (2) David et al. (2014); (3) Davidson et al. (2006); (4) Nagel and Winklbauer (2018); (5) Casini et al. (2012); (6) Ori et al. (2006); (7) Matthews et al. (2008); (8) Muñoz et al. (2006); (9) Ohkawara et al. (2011); (10) Zhang et al. (2016).

### Transmission electron microscopy (TEM)

For the TEM pictures, 4% paraformaldehyde and 2.5% glutaraldehyde in 0.05M cacodylate buffer at pH 7.0 were used to fix stage 11 gastrulae. Bisected samples were rinsed in 0.1M cacodylate and then fix in 0.1M cacodylate containing osmium tetroxide (1%). To visualize the glycocalyx, 1% lanthanum nitrate (Sigma-Aldrich Canada) was added to the fixatives. Samples were washed with 0.1M cacodylate and dehydrated in a series of graded ethanol solutions before embedding in Spurr’s resin. Semi-thin and ultrathin sections were obtained using a Lecia EM UC6 microtome. Sections were stained with 3% uranyl acetate in methanol for 1 hour followed by 10 minutes in Reynold’s lead citrate. Images were taken with a Hitachi HT7700 microscope.

#### Analysis of TEM images

Cell contacts were analysed as previously described (Barua et al. 2021). In short, differences in cell and yolk platelet size were used to identify morphant gastrula tissues in TEM images from 3 embryos from different egg batches for each condition (Barua et al. 2021). Stretches of cell perimeter were treated as contacts when the contours of adjacent cells followed each other. Their abrupt divergence indicated the end of a contact at a non-adhesive gap. Contact angles between cells were measured at the transitions between contacts and interstitial gaps. Contact width was measured as separation distance between membranes, distances were binned in 50 nm wide steps. LSM height in gaps was similarly measured. All contacts or angles in a TEM image were measured; for number of TEM images per treatment (see Figure legends). The error due to random tilting of the sectioning plane relative to the plane of a contact, as estimated from the width variation of tight junctions, can amount to an up to 1.5-fold apparent increase in contact width (Barua et al. 2021). The respective distortion of contact angles broadens the angle distribution by a factor k, by making the narrowest angles appear even narrower and widening the widest angles, depending on the inclination of the sectioning plane (Barua et al. 2017). The factor k depends on the shape and size of the objects sectioned at random and the angle distribution. It is difficult to derive theoretically, given that the gaps in the tissues examined here are a heterogeneous mixture of differently sized bubbles and 3-to-multisided gaps. However, we found that when setting k ≥ 1.5 the corrected maximal and minimal boundary lines would intersect within the observed width range, whereas at a slightly lower k = 1.33 the lines would meet only beyond this range (Fig.S4). Variances between treatments were statistically analyzed using one-way ANOVA. Data visualization and statistical analyses were performed using Graphpad Prism 7 2017 v7.0.3. Vector graphics and figures were assembled using Inkscape v0.92 and v1.0.

## Acknowledgements

We thank A. Chong of the imaging facility at the University of Toronto’s Department of Cell and Systems Biology for help with TEM imaging. Funding was provided to R.W. by the Canadian Institutes of Health Research (PJT-15614) and by the Natural Sciences and Engineering Research Council of Canada (RGPIN-2017-06667).

## Conflict of interests

The authors declare that they have no conflict of interests.

---

**Supplementary Figure S1.**
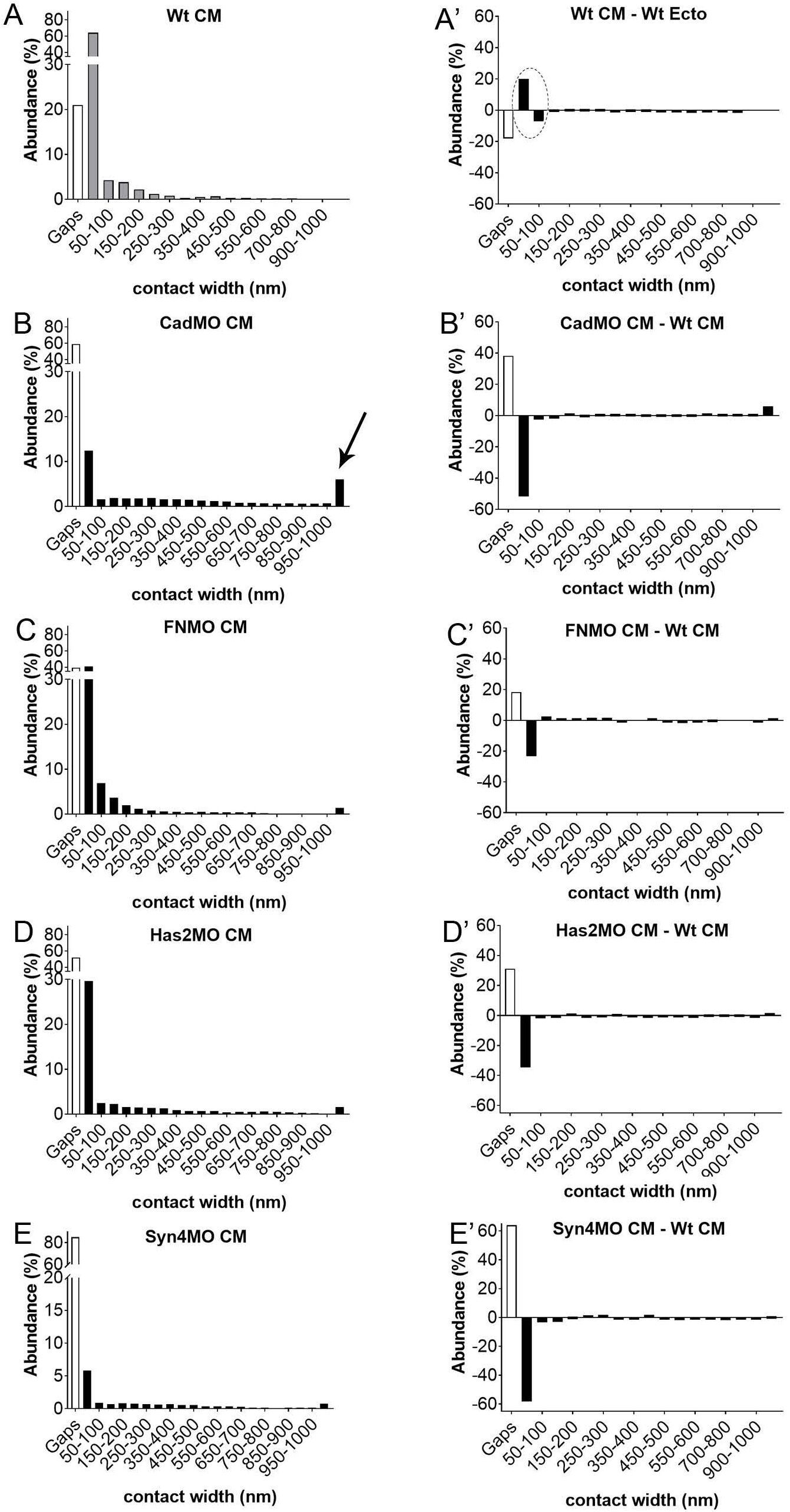
Frequency distributions of contact widths. (A-E) Contact width spectra for normal CM (A), from Barua et al. 2021, and for CM morphants (B-E). Contact widths abundances were collected in 50 nm width bins. (A’) The difference spectrum comparing normal CM and ectoderm shows the signature of glycocalyx I (encircled), suggesting absence of this contact type in the CM. (B’-E’) Difference spectra comparing CM morphants to normal CM (the spectrum for normal CM subtracted bin by bin from respective morphant spectra).

**Supplementary Figure S2.**
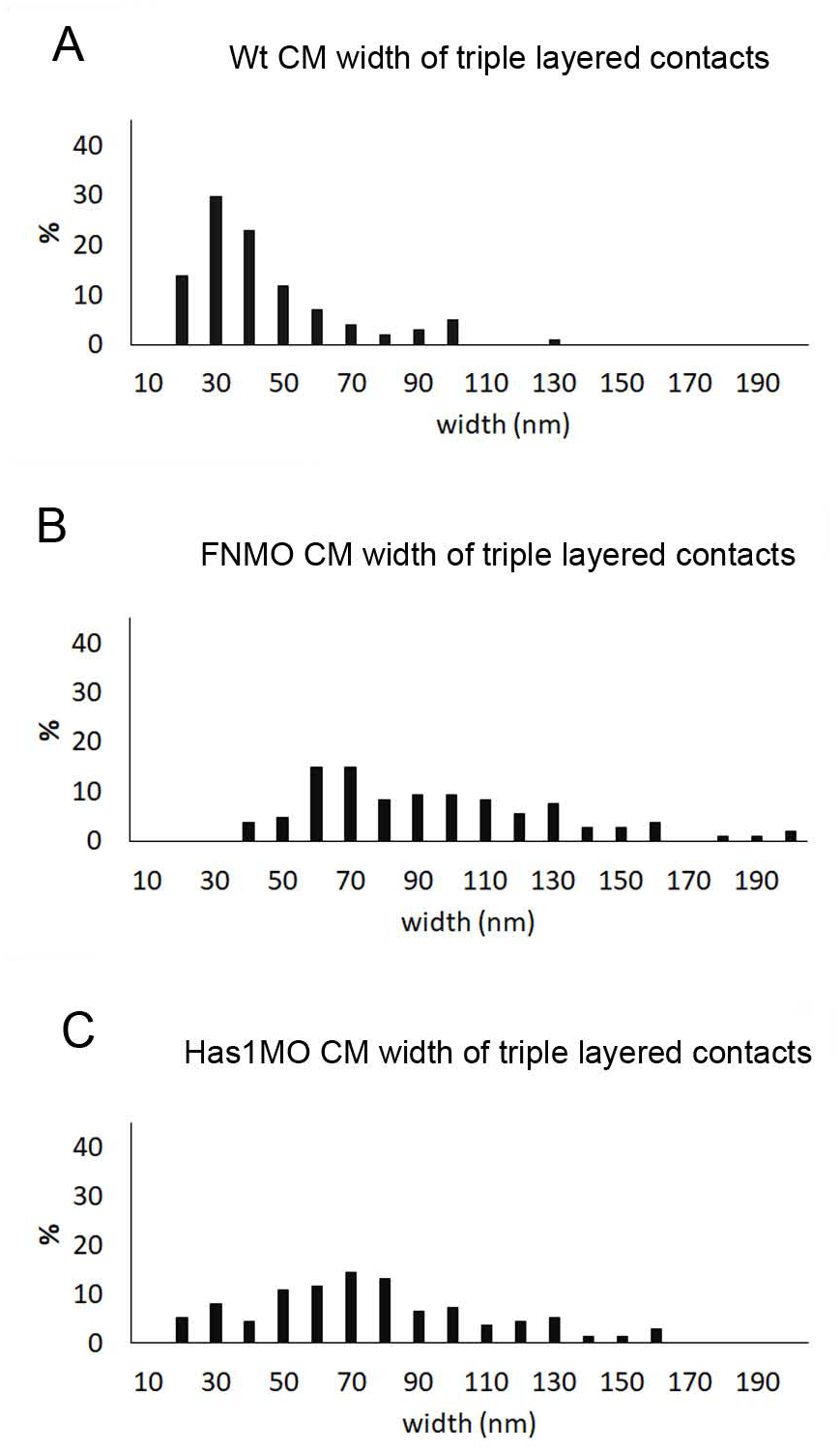
Width frequency distributions of triple-layered (LSM-unlabeled-LSM) contacts. Wt, 101 measurements from 4 TEM images; FN knockdown (FNMO) 107 measurements from 8 TEM images; Has1 knockdown (Has1MO) 138 measurements from 8 TEM images. No triple-layered contacts were seen in C-cad or Syn-4 morphants.

**Supplementary Figure S3.**
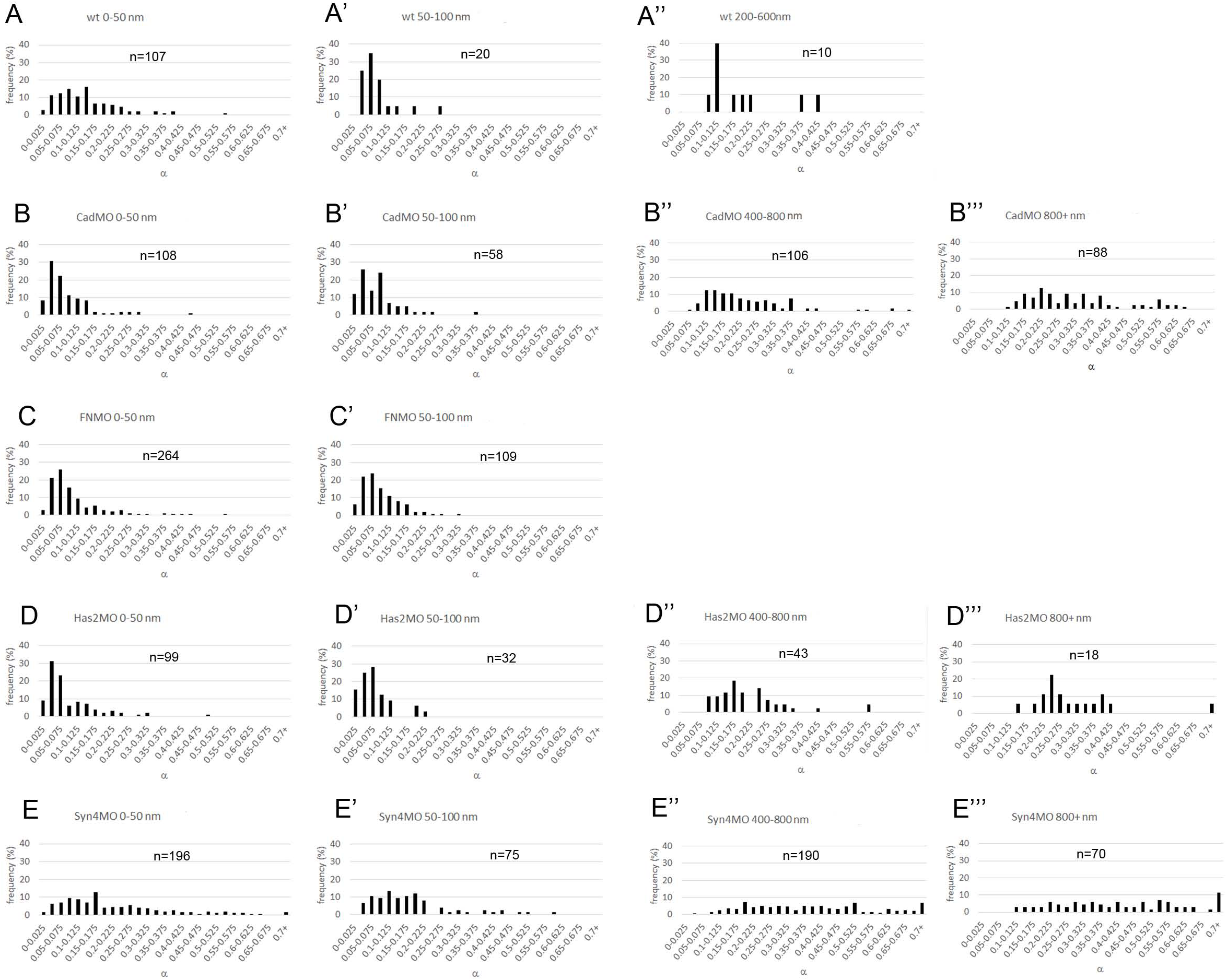
Frequency distributions of α values at different widths. (A-E, A’-E’) Each 0-100 nm width bracket in Fig.8 is broken up into a 0-50 nm and a 50-100 nm bracket, respectively. (A’’-E’’, B’’’-E’’’) Distributions for larger widths brackets not shown in Fig.8. Treatments and width brackets indicated on top of each diagram. n, number of α–w data points.

**Supplementary Figure S4.**
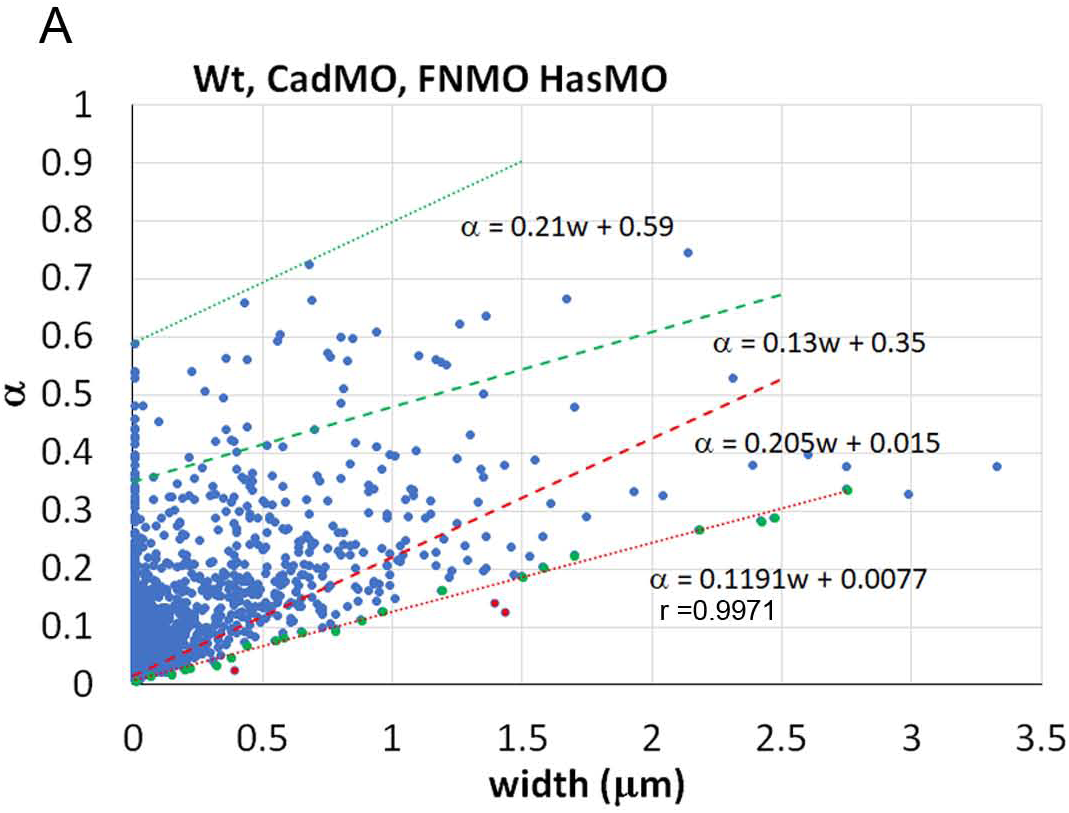
Relative adhesiveness α plotted as a function of contact width w for combined data, to estimate a correction factor k for contact angle values. Dotted lines, regression lines for lowest (red) and highest values (green). Dashed lines, corrections for contact angle distortions due to random orientation of sectioning planes, as described in the Methods section. K = 4/3 is compatible with the intersection of the corrected lines just beyond the α – w distribution. Equations for lines are indicated as in Figure 7.

**Supplementary Figure S5.**
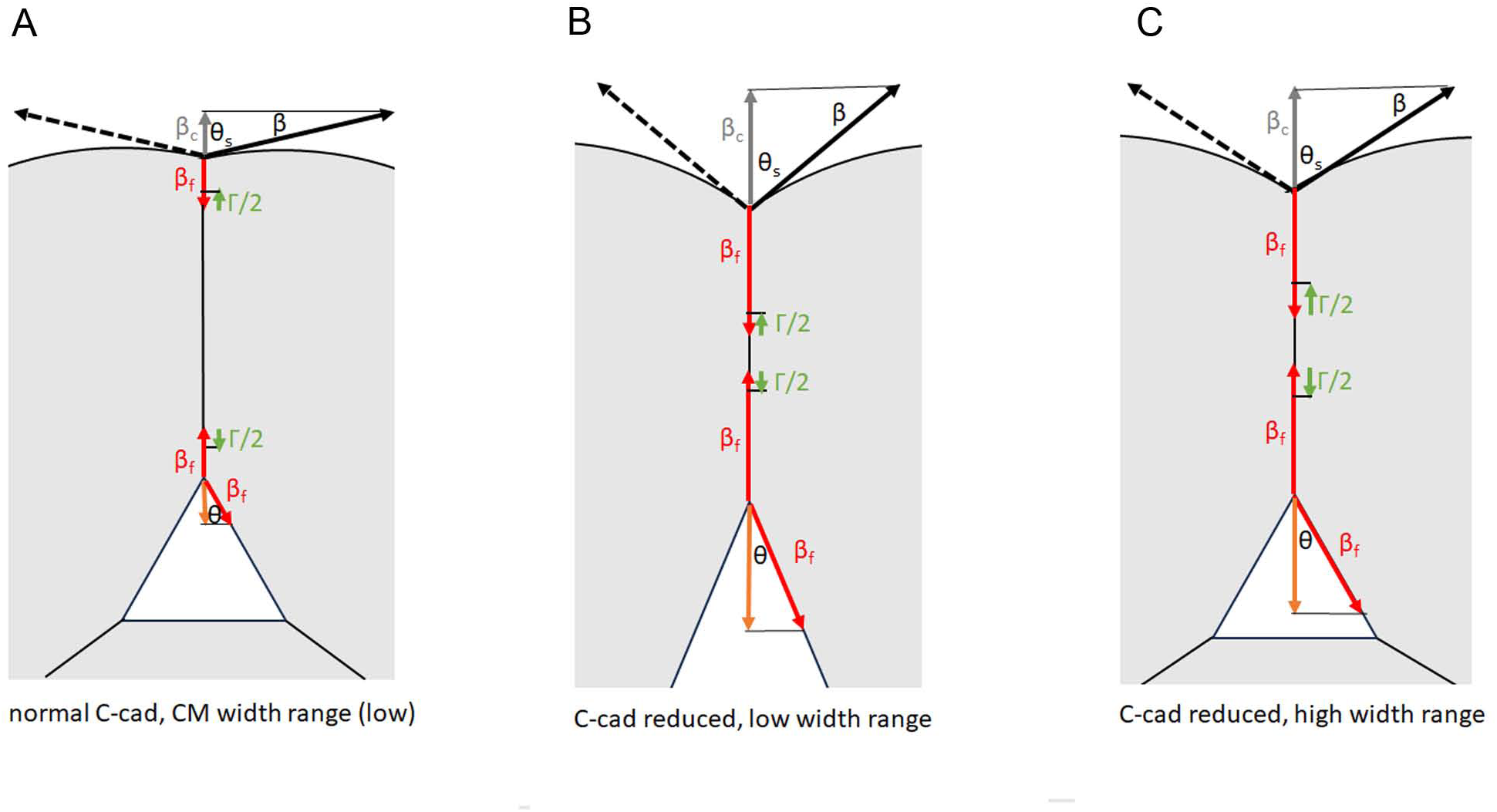
Diagrams schematically depicting the relationships between tensions (shown per cell) and contact angles at tissue surfaces (top) and at interstitial gaps (triangles). (A) In normal CM, cortical tension β at the tissue surface (black) is strongly reduced upon cell adhesion to tension β_f_ (red). Release of binding energy due to adhesion factor interactions at the narrow CM contacts generates an average adhesion tension Γ/2 (green). Tensions β_f_ and Γ/2 balance the resultant tension β_c_ (grey); surface contact angle, θ_s_. The same tensions β_f_ and Γ/2 act at the transition to interstitial gaps, but at the gap surface not β but the much smaller β_f_ balance these tensions (orange), requiring a much smaller contact angle θ. (B) In C-cad depleted tissue, tension β at the free surface remains but it is much less diminished at contacts. In the width range of normal CM, Γ/2 remains the same, contact angle θ must become smaller, and the relative adhesiveness α appears reduced. (C) When the average Γ/2 is increased with contact width much beyond the normal CM range, the contact angle θ at gaps can remain the same or even increase while a lowered angle θ_s_ at the surface still indicates the reduced overall adhesion strength (the difference σ = β – β_c_) due to C-cad depletion.

**Supplementary Table S1.** Relative adhesiveness α at low (w < 250 nm) and high (w ≥ 250 nm) contact widths.

| | Contact width $< 250$ nm | | | | | Contact width $\geq 250$ nm | | | |
| --- | --- | --- | --- | --- | --- | --- | --- | --- | --- |
|  | Av | SD | n | p(MO/wt) |  | Av | SD | n | p (low/high) |
| Wt | 0.1326 | 0.0898 | 143 |  |  | 0.2260 | 0.1380 | 6 | 0.0158 |
| CadMO | 0.0966 | 0.0786 | 259 | 0.0001 |  | 0.2495 | 0.1380 | 249 | 0.0001 |
| FNMO | 0.1030 | 0.0871 | 476 | 0.0004 |  | 0.1853 | 0.1244 | 53 | 0.0001 |
| Has2MO | 0.0842 | 0.0693 | 165 | 0.0001 |  | 0.2040 | 0.1234 | 91 | 0.0001 |
| Syn4MO | 0.2339 | 0.1611 | 423 | 0.0001 |  | 0.3806 | 0.1834 | 408 | 0.0001 |

Av, average of $\alpha$ values; SD, standard deviation; n, number of $\alpha$ -w data points, i.e. gap-contact transitions; p (MO/wt), p-values for significance morphants vs. untreated CM (wt); p (low/high), p-values for significance low vs. high contact widths.

**Supplementary Table S2.**
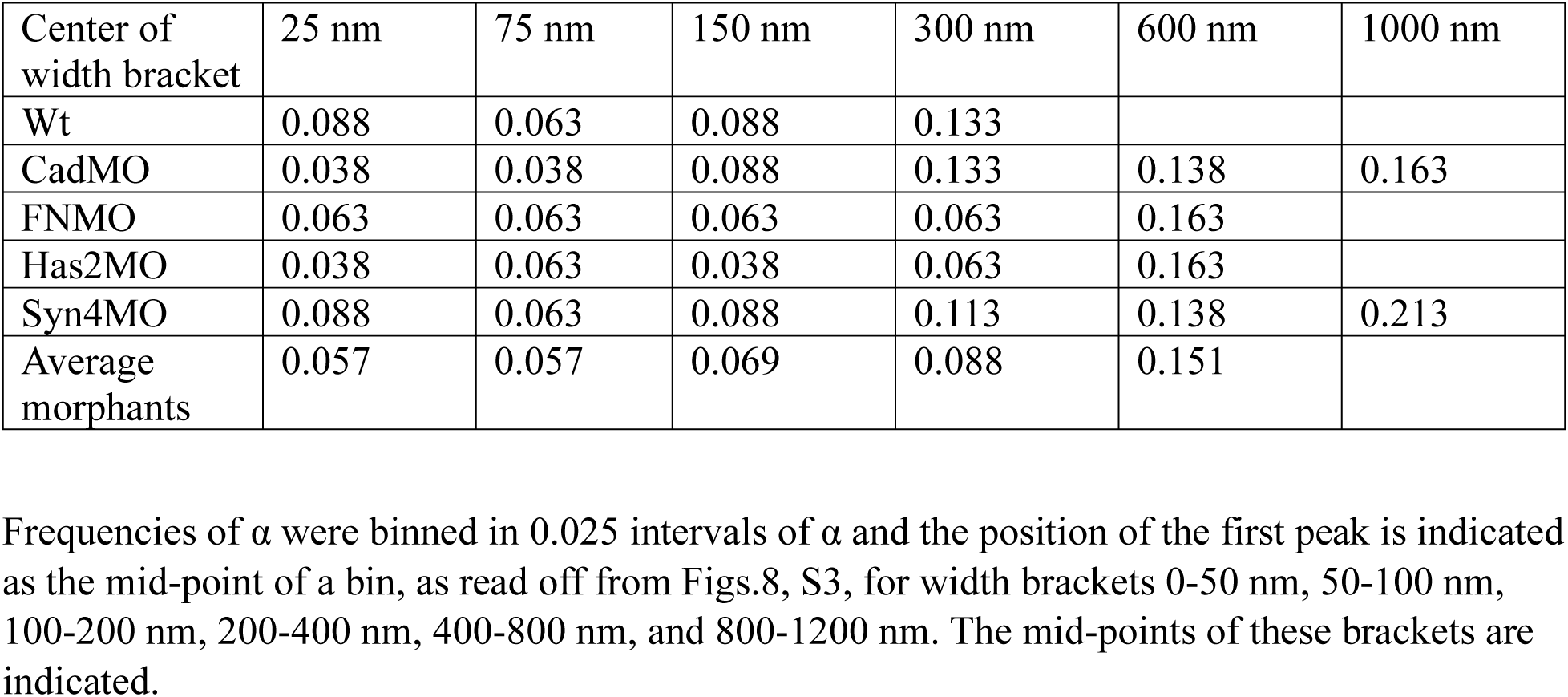
Positions of first peak in α frequency distributions in consecutive width brackets.

**Supplementary Table S3.** Regression line parameters.

| | $\alpha_0$ | $\Delta\alpha/\Delta w$ [$\mu\text{m}^{-1}$] | r | d [nm] |
| --- | --- | --- | --- | --- |
| wt | 0.017 | 0.261 | 0.935 | 130 |
| CadMO | 0.013 | 0.114 | 0.988 | 227 |
| FNMO | 0.014 | 0.119 | 0.978 | 235 |
| HasMO | 0.009 | 0.102 | 0.972 | 172 |
| Syn4MO | 0.023 | 0.074 | 0.960 | 625 |

